# Sex-associated autosomal DNA methylation differences are wide-spread and stable throughout childhood

**DOI:** 10.1101/118265

**Authors:** Matthew Suderman, Andrew Simpkin, Gemma Sharp, Tom Gaunt, Oliver Lyttleton, Wendy McArdle, Susan Ring, George Davey Smith, Caroline Relton

## Abstract

Almost all species show sexual discordance in many traits and diseases. DNA methylation is known to contribute to these differences through well-established mechanisms including X-inactivation in females, imprinting and parent-of-origin effects. Here we investigate sex discordance in DNA methylation throughout childhood in a sample of 700 individuals from the Avon Longitudinal Study of Parents and Children. We show that autosomal sex-discordant methylation is widespread, affecting approximately 12,000 CpG sites at any given age, and stable; at least 8,500 sites are consistently different across all time points and a large proportion discordant in both the fetal and adult brain cortices. Just over 1,000 methylation differences change from birth to late adolescence, 90% of these between birth and around age seven. Sexually discordant CpG sites are enriched in genomic loci containing androgen but not estrogen targets and in genes involved in tissue development but not housekeeping functions. A methylation-derived sex score capturing the variance was calculated at each time point and found to be highly correlated between time points. This score is nominally associated with sex hormone levels in childhood as well as some phenotypes previously linked to sex hormone levels. These findings suggest that sex-discordant autosomal DNA methylation is widespread throughout the genome, likely due to the first androgen exposures *in utero.* It is then stably maintained from birth to late adolescence. Methylation variation at sex-discordant sites within the sexes, as summarized by the methylation sex score, likely reflects *in utero* androgen exposure which is relevant to human health.

**Significance Statement:** Although we know that sex hormones are critical for establishing sexual discordance, less is known about how this discordance is achieved and maintained. Here we present evidence for widespread differences in DNA methylation between male and female children. We show that most of these differences are established prenatally, likely due to the first androgen exposures *in utero,* and then stably maintained throughout childhood, despite extreme fluctuations in the levels of these very same hormones. Our results support a role for DNA methylation as a means for recording and maintaining the effects of exposure to sex hormones and thus to better understand sexual variation and how it is driven by the prenatal environment.

## Introduction

Although males and females appear to be quite similar, being composed of identical cell types, tissues and organs, there are fundamental differences between them (1). GenderMedDB (http://gendermeddb.charite.de/) lists over 11,000 publications describing the sex-specificity of a wide range of diseases affecting everything from immune response (2) to mental health (3). Given the close relationship between environmental exposures and disease, it is not surprising that many exposures also elicit sex-specific responses (4).

These sex-specific exposure-disease relationships are likely due to molecular differences. The most obvious difference is chromosomal, with males having a copy of the paternal Y chromosome rather than the X chromosome. The presence of a Y chromosome or multiple X chromosomes alone is sufficient to induce changes in gene regulation and expression (5, 6). The stronger female immune response is likely stems from the fact that an immune-enriched 15% of genes on the X chromosome escape X-inactivation resulting in double-dosage of these genes (7). Even in the presence of a Y chromosome, male development can be disrupted by a malfunctioning *SRY* gene or downstream *SOX9* gene and lead to female development (8). Male sex hormone levels are also known to have an effect. These hormones are typically quite high in males during gestation, drop shortly before birth and then surge shortly after birth, generating many of the lasting male characteristics in the brain (9). Exposure of rodent female pups to slightly higher levels *in utero* leads to enhanced male phenotypes, such as a larger sexually dimorphic nucleus induced only by closer proximity to other developing males (10) and enhanced male patterns of aromatase-expressing neurons in the brain (11). Conversely, hormone reduction due to gonadectomy or removal of the pituitary gland removes nearly all male-specific gene expression in the mouse liver (12, 13). However, male-specific expression can be mostly restored by growth hormone treatment (13).

Male and female developmental trajectories are manifest in life-long sex-specific gene expression patterns (14) that are also highly tissue-specific and appear to be evolutionarily conserved. In the mouse liver, as many as 70% of genes have sexually dimorphic gene expression patterns compared to only 14% in the brain (15). Sexually dimorphic gene expression in the occipital cortex is highly conserved across several primates (16).

Sex-specific gene expression patterns may be mediated by a number of different mechanisms including DNA methylation (17). In fact, several studies indicate that DNA methylation levels can be modified by sex hormones. For example, injection of testosterone into female mice at birth induces male-associated methylation patterns in the bed nucleus of the stria terminalis (18). Fluctuations in estradiol levels and exposure to estradiol-imitating compounds like Bisphenol A (BPA) have also been linked to DNA methylation changes (19-21). The speed of these methylation changes suggests that sex hormones may interact directly with biological pathways responsible for creating and maintaining methylation marks (for a review, see (22)).

Indeed, several human studies identify sex-specific methylation differences at candidate autosomal loci (23-26), globally (27-29) and at multiple loci across the genome using microarrays in a variety of cell types including saliva (30, 31), cord blood (32), peripheral blood (33-37), prefrontal cortex (31, 38, 39) and colon (40). Although a large proportion of these reported differences may be erroneous due to microarray probes targeting DNA sequences found on both autosomes and sex chromosomes (35, 41), removal of these ‘cross-reactive’ probes from analyses still identifies hundreds of autosomal sex-specific DNA methylation differences (32, 35, 37-39).

Lacking in this literature is an investigation of how sex-specific DNA methylation changes throughout life in response to development and sex hormone fluctuations. Here we investigate how sex-specific DNA methylation changes throughout childhood, from birth to 17 years of age, including puberty, one of the most dramatic changes in sex hormone levels. We analyze over 2000 DNA methylation profiles from blood DNA obtained using the Illumina Infinium^®^ HumanMethylation 450K BeadChip assay. The profiles were derived from cord blood at birth and peripheral blood in childhood (around 7 years-old) and adolescence (around 15-17 years-old) from children enrolled in the Avon Longitudinal Study of Parents and Children (ALSPAC) (42, 43) (Fig 1 and Table 1). The DNA methylation profiles form part of the Accessible Resource for Integrated Epigenomics Studies (ARIES) dataset (44).

**Fig 1.**
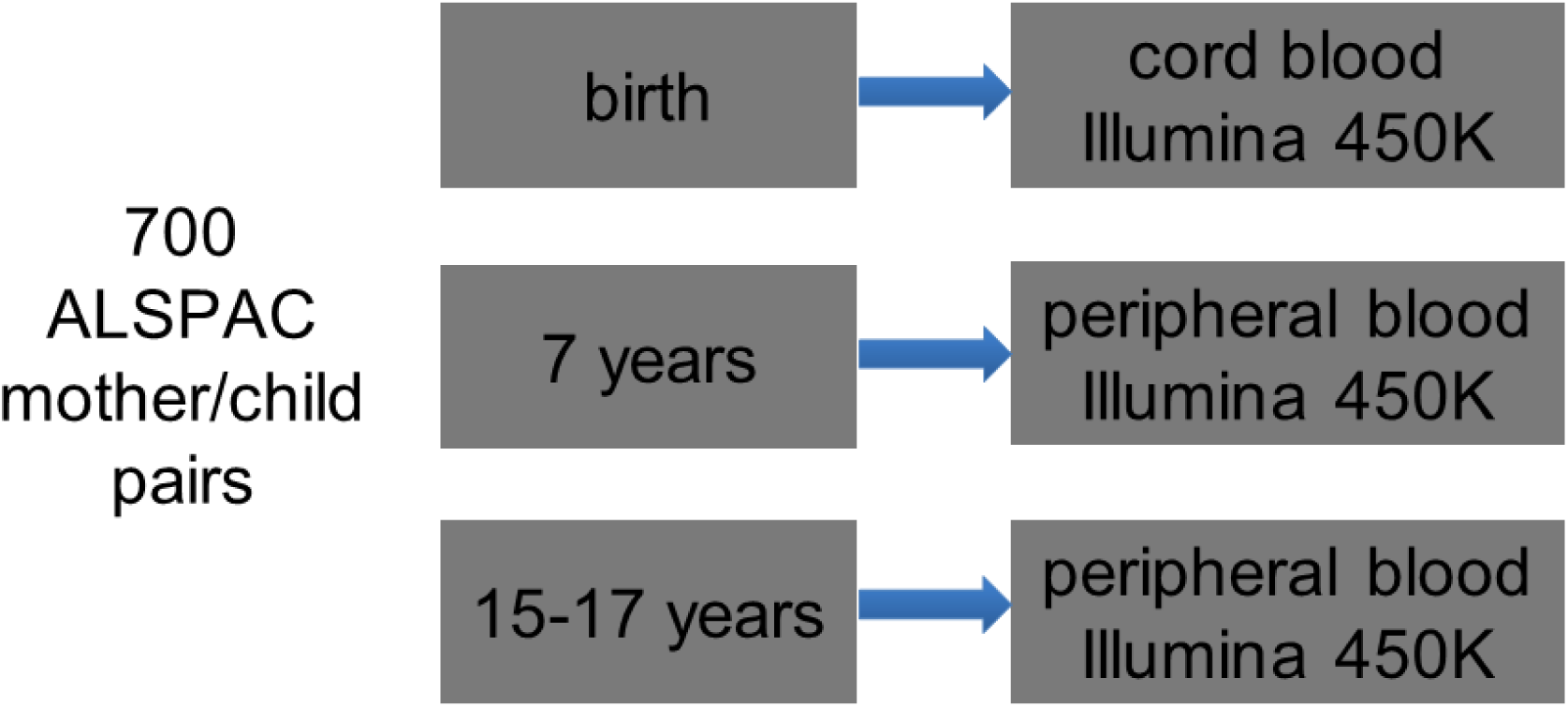
Accessible Resource for Integrated Epigenomics Studies (ARIES) dataset.

**Table 1.** Characteristics of the ARIES sample by sex. Characteristics of the ARIES sample by sex. T-tests are used to test numeric differences between males and females, and Pearson’s chi-square tests to test differences in proportions. Data was collected for most individuals at birth, age 7, age 15-17. Characteristics of the sample for each time point are given in Table S1.

| variables | males |  | females |  | p-value |
| --- | --- | --- | --- | --- | --- |
|  | mean/n | sd/% | mean | sd/% |  |
| n = 700 | 335 |  | 365 |  |  |
| gestational age | 39.5 | 1.5 | 39.7 | 1.5 | 0.07704 |
| birthweight | 3569.1 | 502.9 | 3409.8 | 459.8 | 0.00002 |
| mother age (months) | 364.9 | 54.6 | 355.1 | 52.4 | 0.01608 |
| parity | 0.8 | 0.8 | 0.7 | 0.8 | 0.37894 |
| prenatal smoking |  |  |  |  |  |
| current | 47 | 14.0 | 51 | 14.0 | 1.00000 |
| quit | 15 | 4.5 | 27 | 7.4 | 0.14277 |
| never | 248 | 74.0 | 259 | 71.0 | 0.41017 |
| caesarean | 29 | 8.7 | 32 | 8.8 | 1.00000 |
| smoking daily (15-17y) | 35 | 10.4 | 44 | 12.1 | 0.58116 |

## Results

### DNA methylation at thousands of autosomal CpG sites remains stably sex-specific over time

Between 11-13,000 CpG sites are differentially methylated between males and females at each of the three time points: birth, childhood, adolescence (Bonferroni-adjusted p < 0.05 and >1% methylation difference; Spreadsheet S1). In total, 17,083 CpG sites are differentially methylated in at least one time point. A remarkable 85% of these sites are more methylated in males than females in each time point.

Each pair of time points agrees on 70-80% of differentially methylated CpG sites, with 8,509 sites differentially methylated between the sexes at all three time points (Fig 2). At each of these 8,509 sites, direction of association is also conserved. At almost half of these sites, methylation levels in both males and females show little evidence of change (4,056 or 47.7% of 8,509 sites have coefficients of age in multilevel regression models roughly equal to 0, unadjusted p > 0.05; see Supplementary Materials and Methods). For a small subset (315 or 3.7% of 8,509 sites), methylation levels do appear to have changed between time points (i.e. at least one change statistic has Bonferroni adjusted p-value < 0.05), but sex-associated methylation differences remain.

**Fig 2.**
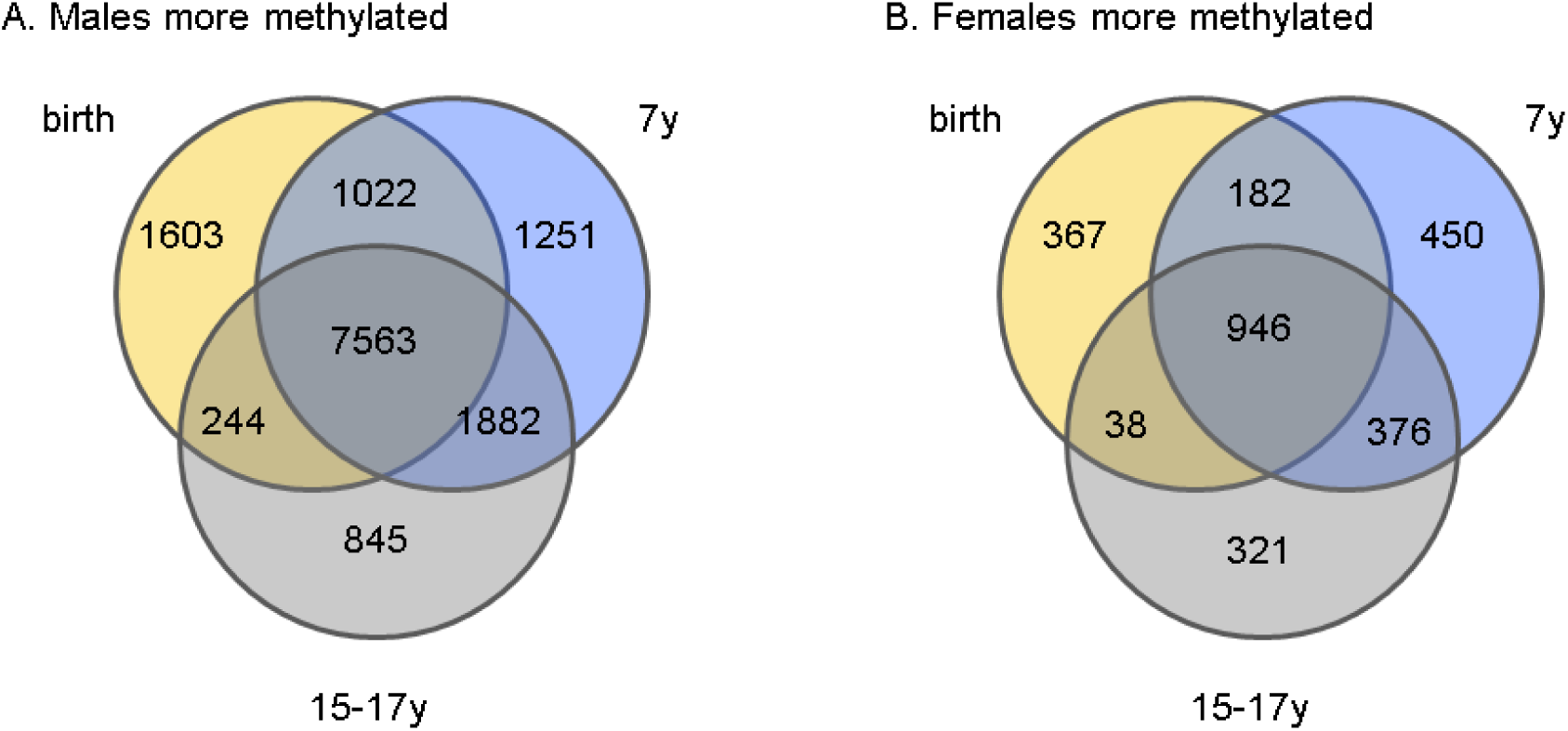
Numbers of sex-specific CpG sites at each of the time points: birth, 7y and 15-17y. Numbers of sites more methylated in males (A) and in females (B) are shown.

### Genes more methylated in males and females enriched for different biological processes

CpG sites more methylated in males are near genes enriched for developmental Gene Ontology biological processes (45) (Spreadsheet S2). These include processes involving post-embryonic morphogenesis and development of bones, heart and skin as well as differentiation of osteoclasts, neurons and cardioblasts. Consistent with overall higher DNA methylation in males, RNA PolII transcription activators are more highly methylated in males, suggesting gene expression levels may be higher in females. However, in apparent contradiction with this prediction, genes involved in DNA methylation are also consistently more methylated in males including: *ATF7IP, BAZ2A, DNMT3A, DNMT3B, GNAS, KDM1B, PLD6, SPI1, TET1, UHRF1.* Many of these are also involved in gene imprinting as well, including *EED, IGF2, KCNQ1, NDN, PEG3* and *ZIM2.* Genes related to female pregnancy and placenta development are more methylated in males, consistent with them being less active in males.

CpG sites more methylated in females were enriched near genes that respond to toxic exposures, possibly playing a role in the known sexual dimorphic responses to such exposures (46-49). Genes *CYP1A1* and *TH* are known to respond to herbicide exposure (50, 51). Two of the three CpG sites near *CYP1A1* that are more methylated in females are similarly more methylated in the cord blood of infants with prenatal tobacco exposure (52). Another site near *GFI1* that is less methylated in response to prenatal tobacco is also consistently less methylated in females. Female methylated genes *TH* and *NTRK1* are known to respond to nicotine (53, 54). Although we are not aware that these genes are differentially methylated between males and females in response to toxic exposures, the fact that they are differentially methylated between the sexes and in response to exposure is consistent with a sexually dimorphic response to toxic exposures.

Other CpG sites more methylated in females are enriched near genes involved in reproduction. Four such sites appear near the transcription start site of *PIWIL2,* a gene known to play roles in spermatogenesis and oogenesis and found differentially methylated between infertile and control males (55). Five of the CpG sites are just upstream of *PRDM9,* a gene involved in determining meiotic recombination hotspots in both humans and mice (56). Four of the CpG sites are within the first intron of lincRNA *NBR2,* just upstream of *BRCA1,* a gene activated by sex hormones estrogen and progesterone (57).

Few housekeeping genes (58) are linked to sex-specific DNA methylation. This is expected since the products of the genes would be needed by both males and females. Specifically, 1440 housekeeping genes were differentially methylated, whereas 1728 were expected by chance (p < 2×10^−15^, Fisher’s exact test). Given that housekeeping genes tend to be proximal to CpG islands, this would appear to suggest that sex-specific DNA methylation would occur in low CpG frequency regions. We however find the opposite, the median CpG frequency at sex-associated CpG sites is 0.6 compared to 0.45 at other CpG sites (p < 2×10^−16^, Wilcoxon rank sum test). CpG frequency was measured as the ‘normalized CpG frequency’ (59) for the 200bp sequence centered at the CpG site.

### Genes involved in sexual development are enriched for sex-specific DNA methylation

Given the enrichment of developmental processes, we asked whether any genes with sex-specific DNA methylation had been linked to sexual development specifically. Table S2 lists genes that have been linked to specific components of sexual development.

In most cases, these genes are enriched for sex-specific methylation as indicated by low enrichment p-values in Table S2. For example, a CpG site in the second intron of *SOX9* is consistently about 3% more methylated in males than females (cg13058710). A CpG site upstream (cg15265222) and another downstream (cg03688324) of the first exon of *FGF9* are both more methylated in adolescent females. All but one of the genes linked to external genitalia development (*ZFPM2*) is differentially methylated between males and females, and all but two (*ATF3* and *SSH*) of the remaining are more methylated in males than females at each time point. *ATF3* is more methylated in males only in cord blood, and *SHH* has sites more methylated in either males or females at each time point. Almost all of 16 autosomal HOX genes linked to sexual development (*EMX2, HOXA11, HOXA13, HOXA7, IRX3, LBX2, LHX1, LHX9, MKX* and *TGIF1*) have greater methylation in males at each ARIES time point. The two exceptions, *LHX8* and *PBX4,* are both more methylated in females at each time point. *LHX8* is involved in oogenesis and *PBX4* in spermatogenesis.

### Genes with sex-specific DNA methylation are enriched for androgen but only weakly for estrogen targets

Given the fundamental roles of sex hormones in sexual development and in maintaining sex differences, we investigated the extent to which genes with sex-specific methylation are targeted by sex hormones or hormone receptors.

We first considered testosterone, an androgen that plays a key role in the development of male characteristics. Testosterone targets were defined as genes differentially methylated in female mice after being injected with testosterone at birth (18). Of the genes differentially methylated in the mouse striatum, 975 have human homologs and >450 are differentially methylated at each time point in our study (Table 2). Of those differentially methylated in the mouse bed nucleus of the stria terminalis (BNST), 497 have human homologs and >240 are differentially methylated at each time point in our study (p < 1.8×10^−11^).

**Table 2.** Overlap of sex hormone targets with sex-specific DNA methylation. Just over 4000 genes with sex-specific methylation in humans (ARIES) have homologs in mouse. These include about 50% of the 975 genes targeted by testosterone in mouse striatum (18) and less than 40% of the 632 genes targeted by estrogen in mouse hippocampus (62).

| ARIES |  | Testosterone targets in mouse striatum |  |  | Estrogen targets in the mouse hippocampus |  |  |
| --- | --- | --- | --- | --- | --- | --- | --- |
| Time point | Sex-specific genes | Genes | Intersect | P-value | Genes | Intersect | P-value |
| cord | 4761 | 975 | 454 | $1.4 \times 10^{-12}$ | 632 | 250 | 0.023 |
| 7y | 4960 | | 488 | $1.7 \times 10^{-17}$ | | 253 | 0.075 |
| 15-17y | 4585 | | 466 | $2.7 \times 10^{-19}$ | | 239 | 0.037 |

Testosterone and other androgens control gene expression through binding to the androgen receptor. Androgen receptor targets were identified in a previous study of dihydrotestosterone stimulated prostate cancer cell lines (60). Of the 193 bound genes, 156 have DNA methylation measurements in our study and 51, 61 and 55 are differentially methylated at birth, childhood and adolescence, respectively, in our study (unadjusted p = 0.044, 0.0012 and 0.0042, respectively, Fisher’s exact test).

Estrogens are also likely candidates for driving sex-specific DNA methylation since they play a key role in the development and regulation of the female reproductive system and are known to affect DNA methylation levels directly (22). Estrogen targets were defined in three different ways: estrogen binding, methylation changes following estradiol treatment, and methylation and expression changes following Bisphenol A treatment.

Estrogen binding sites were identified from a previous study of estrogen receptor binding in two different estrogen-responsive human cell lines: ECC-1 (endometrial cancer) and T47d (breast cancer) (61). In the study, cells were exposed to BPA, genistein (found in soybean), or 17β-estradiol (an endogenous estrogen) in order to induce greater estrogen receptor activity. There was a weak enrichment at each ARIES time point only for the CpG sites coinciding with estrogen binding stimulated by 17β-estradiol in each of the cell types (for birth, childhood and adolescence, there were ~20 such CpG sites for unadjusted p = 0.04, 0.10 and 0.08, respectively; Fisher’s exact test).

Estrogen targets were also defined as genes differentially methylated in a previous study of hippocampal DNA methylation in female ovariectomized mice with and without estradiol treatment (62). We observed slight enrichment of these targets among genes differentially methylated at birth (unadjusted p = 0.023 for 185 of 632 human homologs, Fisher’s exact test) and at adolescence (unadjusted p = 0.037 for 239 of 632 human homologs) (Table 2).

Estrogen targets were finally defined as genes differentially expressed and methylated in normal-like human breast epithelial cell line MCF-10F cells following treatment with estradiol-imitating compound Bisphenol A (BPA) (19). We observe enrichment of genes with lower expression and higher methylation among those differentially methylated in our study at each time point. Of the genes responding to low BPA treatment, 18-19 are differentially methylated at each ARIES time point (unadjusted p = 0.00009, 0.0048 and 0.0018 for birth, childhood and adolescence, respectively; Fisher’s exact test). Of those responding to high BPA treatment, 35-41 genes are differentially methylated (unadjusted p = 0.0024, 0.074 and 0.04 for birth, childhood and adolescence, respectively). There were a small number of genes with higher expression and lower methylation with almost no overlap with sex-specific methylation in ARIES.

### Genes with sex-specific methylation tend to be highly expressed in the adrenal cortex

We asked which cell type expression pattern, among 79 different cell types (63), was best represented by the set of genes differentially methylated between the sexes at each time point in ARIES. In particular, each gene was identified as highly expressed in a cell type if the expression microarray probe intensity was in the upper quartile for that cell type. The strongest overlap we observed was between adrenal cortex expression and sex-specific genes at 7 years of age in ARIES (Bonferroni adjusted p = 1.04×10^−4^, Fisher’s exact test). The other time points have weaker enrichments with adrenal cortex expression (p = 0.012 for 17y and p = 0.062 for cord). There are similar but weaker enrichments for liver expression and sex-specific genes at age 17y (p = 0.05) and ovary expression and sex-specific genes at age 7y (p = 0.05). In spite of the fact that our methylation data was derived from blood, there appears to be no enrichment for whole blood gene expression in methylation differences at any time point (p > 0.9).

### Sex-specific CpG sites consistently enriched in imprinted regions

Imprinted genes are genes expressed from only the allele inherited from one parent. Imprinting control regions (ICRs) are genomic regions that exert control over the imprinting of nearby genes typically using DNA methylation as a silencing mechanism. For many genes, imprinting is sex-specific. For example, a recent study identified 347 autosomal genes with sex-specific imprinting features in the mouse brain (64).

We found that imprinting control regions (65) are enriched with sex-specific CpG sites (p < 10^−22^, Fisher’s exact test) as are regions within 10Kb of an imprinted gene (p < 10^−15^). We did not however observe any enrichment around human homologs of the 347 autosomal mouse genes with sex-specific imprinting features, in spite of the fact that these genes are enriched for testosterone targets (see Supplementary Text S1).

### Males more variably methylated than females

Given the apparent dependence of male methylation patterns on sex hormones and the fact that testosterone levels are known to vary significantly between males, we reasoned that methylation variance might be higher in males. To test this we applied double generalized linear models in order to simultaneously test mean and variance differences between males and females (66). Similarly to methylation mean differences, methylation variance differences are strongly conserved over time and, as expected, males tend to have more variable methylation; about 80% of sites with mean and variance differences have greater variance in males (Table 3).

**Table 3.** Sex-specific methylation variation. CpG sites with sex-specific methylation levels and variance. The only overlaps that survive adjustment for multiple tests (Bonferroni adjusted p < 0.05; Fisher’s exact test) are within each sex, between different ARIES time points. For each, p < 10^−50^ (Fisher’s exact test).

|  |  | Males more variable |  |  | Females more variable |  |  |
| --- | --- | --- | --- | --- | --- | --- | --- |
|  |  | birth | 7y | 15-17y | birth | 7y | 15-17y |
| Males more variable | birth | 652 | 456 | 331 | 0 | 4 | 4 |
|  | 7y | 456 | 1579 | 581 | 24 | 0 | 6 |
|  | 15-17y | 331 | 581 | 785 | 10 | 1 | 0 |
| Females more variable | birth | 0 | 24 | 10 | 206 | 92 | 55 |
|  | 7y | 4 | 0 | 1 | 92 | 299 | 83 |
|  | 15-17y | 4 | 6 | 0 | 55 | 83 | 154 |

Since methylation variance is often associated with methylation levels, we asked if the difference in variance is simply a function of males have generally higher methylation levels than females. We tested this in two ways. First, we restricted analyses to CpG sites with at most a 2% methylation difference between males and females reasoning that such a small mean difference would be unlikely to induce a measurable variance difference. For each time point, this restricted set of differentially methylated sites included 10 or fewer sites with greater variance in females and more than 200 sites with greater variance in males. Second, we performed random selections of CpG sites more methylated in males and random selections of CpG sites more methylated in females such that the distributions of the mean methylation levels of each set in each sex was identical (see Supplementary Materials and Methods). Across all random selections, males were 3 times (median) more likely in cord blood to have higher variance. At age 7, males were more than 4 times and at age 17 more than 1.5 times more likely to have higher variance. We conclude that higher variance in males is due to something other than having different mean methylation levels.

### Methylation-derived sex score variation preserved over time

Given the variability of sex-specific traits within the sexes, we asked if DNA methylation could be used as a proxy for this variability. Methylation variation was therefore summarized as the weighted mean of methylation at sex-specific CpG sites. Coefficients from the regression models used to identify sex-specific methylation were used as weights (see Supplementary Materials and Methods). As expected, the resulting scores appear as a mixture of two different distributions, male and female with considerable within-sex variation (Fig 3). Using a standard expectation maximization algorithm to fit a bimodal normal mixture to the score, we found that posterior probabilities of membership in each distribution corresponded almost exactly to sex designations. For cord blood, 7y peripheral blood and 17y peripheral blood there were only 11, 13 and 18 misclassifications, respectively. In each case, the misclassification occurred at a single time point with a correct classification at the other time points. Posterior probabilities of misclassifications were not clustered near 0.5 indicating that any were close calls.

**Fig 3.**
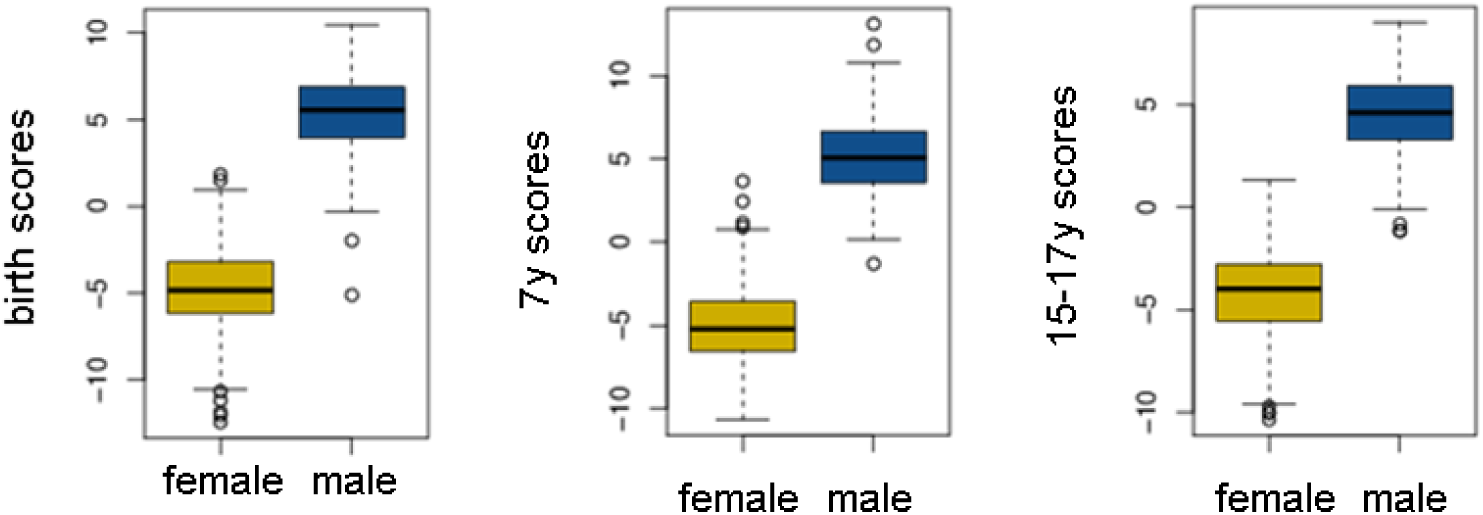
Sex scores for each time point, each sex shown separately.

Within each sex, sex scores were generally correlated between time points, particularly between age 7y and 15-17y but not between birth and 15-17y in males (Fig 4). For some reason this correlation is extremely weak in males compared to females (Spearman correlation R = 0.2 in females compared to R = 0.016 in males, p = 0.01).

**Fig 4.**
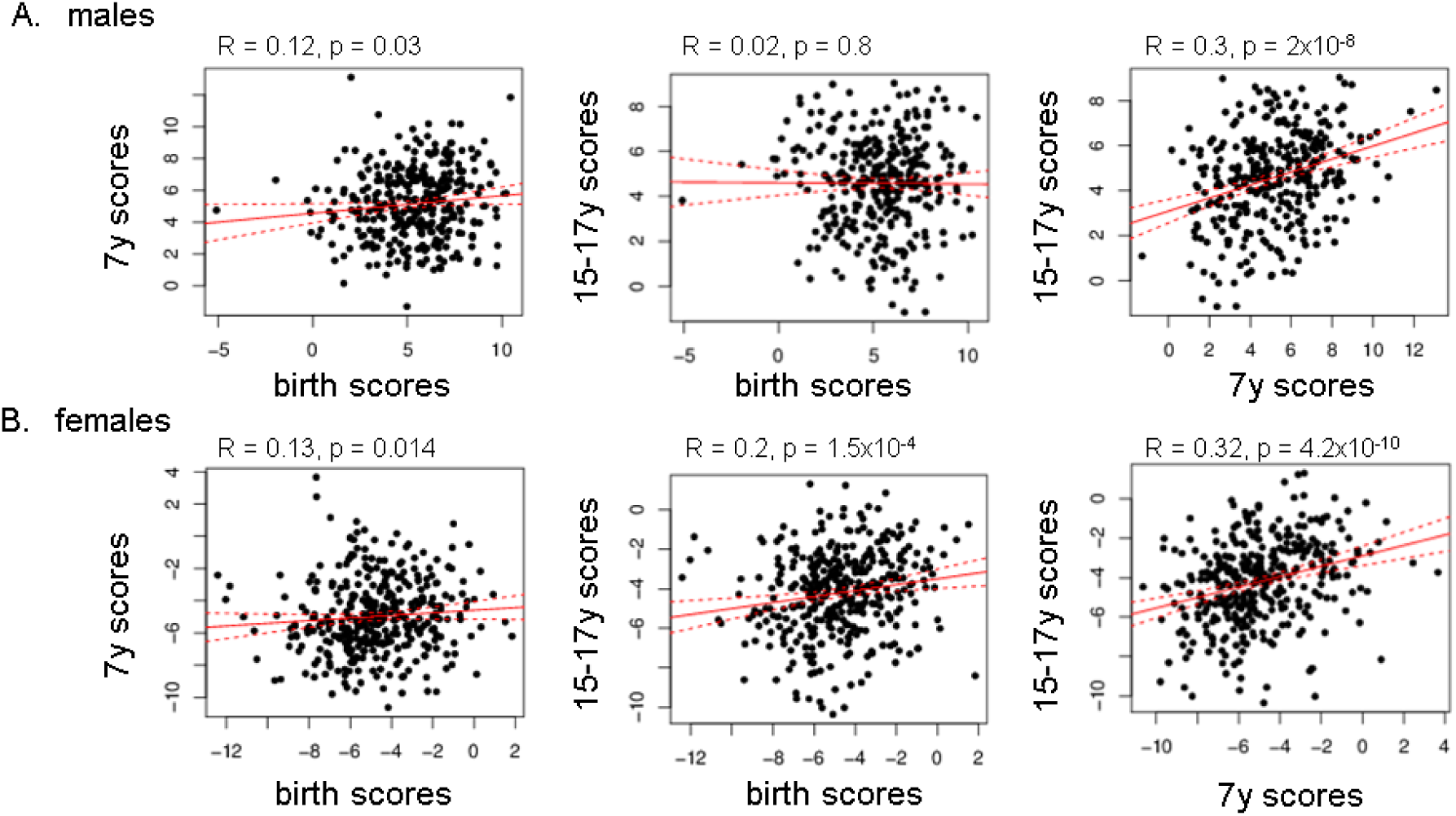
Comparison of sex scores between time points. Solid line depicts trend line. Dashed lines above and below depict the 95% confidence interval. Correlation coefficient is Spearman’s Rho.

This difference of preservation of male and female sex scores may be partially due to the fact that methylation at CpG sites differentially methylated between males and females are more highly correlated over time in females than in males (Table S3). The differences are not large but have very low p-values (p < 1.2×10^−11^).

### Methylation sex scores are associated with estimated cell type ratios and weakly with sex hormone levels

Given the potential influence of hormones on sex-specific DNA methylation, we asked whether or not the sex score is associated with sex hormone-related molecules available in ALSPAC: testosterone, androstenedione, dehydroepiandrosterone (DHEAS), growth hormone binding protein (GHBP), and sex hormone binding globulin (SHBG). Testosterone and SHBG were measured in males at ages 9.8, 11.7, 13.8, 15.4 and 17.7 years of age (67). Testosterone was measured at age 8.5 years in females only. Androstenedione, DHEAS, GHBP and SHBG were measured in males and females at age 8.5 years. Measurements were performed only for about 20% of the individuals in ARIES so power to detect associations is low. Indeed, no associations survived adjustment for multiple tests (at Bonferroni adjusted p < 0.05). Only two were nominally associated in females: GHBP and SHBG at 8.5y with the sex score at 15-17y (Spearman’s rho = -0.38, 0.34; p = 0.003, 0.005), and four were nominally associated in males: testosterone at 13.8y with the 7y sex score (Spearman’s rho = 0.25, p = 0.025), testosterone at 11.7y with the cord sex score (Spearman’s rho = 0.22, p = 0.05), SHBG at 13.8y with the 15-17y sex score (Spearman’s rho = -0.23, p = 0.035), and SHBG at 15.4y with the 15-17y sex score (Spearman’s rho = -0.23, p = 0.034).

Evidence for a causal relationship of sex hormone levels on DNA methylation was evaluated within a two-sample Mendelian randomization framework (68) using SNP-sex hormone associations from published genome-wide association studies (GWAS) and SNP-sex score associations derived from ARIES. Common SNPs associated with sex hormones were obtained from the GWAS catalogue (69). Twenty-one SNPs were identified by filtering the catalogue study descriptions for “testosterone”, “sex hormone-binding globulin”, “SHBG” or “DHT”. Sixteen of these SNPs or their tagging SNPs were measured in ARIES. Association statistics from the sex hormone and sex score associations were used to calculate a Wald ratio estimate for the causal relationship of sex hormones on the sex score. Only estimates for testosterone-associated SNP rs4149056 (70) with the sex score at age 7y was nominally significant at p < 0.05(70) (Table S4).

### Methylation sex scores are not strongly associated with exposures and phenotypes linked to male sex hormones

In rats, intrauterine proximity of a female fetus to male fetuses is known to increase exposure of the fetus to higher testosterone levels (10). Analogously in humans, it may be possible for the hormonal environment of a fetus to be affected by the sexes of older siblings. In 7 year old females, the number of older brothers compared to older sisters is weakly negatively associated with sex score (Spearman rho = -0.12, p=0.067, n = 285), and in 7 and 15-17 year old males, having a brother as the youngest older sibling causes a slight increase in sex score, i.e. methylation patterns are more masculine (mean sex score increase ~ 0.7, n ~ 140, p = 0.08, Wilcoxon rank-sum test). Other tests showed now evidence of association (p > 0.2).

Hand grip strength is highly sexually dimorphic (e.g. (71)) and can be improved by testosterone therapy in men with low testosterone levels (72, 73). In ALSPAC, there are associations between grip strength measured at 11.5 years of age and hormones measured at age 8.5 years of age in both males and females (unadjusted p = 0.00015-0.05, Table S5). Grip strength and sex score were positively associated in males at age 15-17y (Spearman’s rho = 0.13, p = 0.026, n=377) and possibly negatively associated with females at the same age (Spearman’s rho = -0.07, p = 0.2, n=406).

There is evidence that the 2D:4D finger ratio, the ratio of the lengths of the index (2nd) and ring (4th) fingers, may be affected by prenatal testosterone exposure, although the relationship at best appears to be complex (74). A more reliable indicator is anogenital distance (75). Unfortunately anogenital distance is not available in ALSPAC. For 2D:4D ratio, there is a highly strong decrease in males ratios compared to females (p = 10^−36^, Wilcoxon rank-sum test). However, we do not observe any association with sex score apart from a very weak negative association in females between the right hand 2D:4D ratio and sex score at age 15-17y (Spearman’s Rho = -0.1, p = 0.073, n=423).

Finally, we tested associations between sex score and two different scales for gender role behaviour, the Pre-School Activities Inventory (PSAI) measured at 42 months and its adapted and shortened version called the Children’s Activities Inventory (CAI) measured at 8.5 years of age. Although PSAI and CAI are highly different between males and females, there appear to be no associations within individual sexes with sex score.

### One in 15 methylation differences changes over time

Although most sex differences remain unaltered from birth to 15-17 years of age, the differences at a small percentage of CpG sites do change over time. Fig 5 depicts two examples, one for which the methylation difference shrinks and another for which it grows followed by four examples that neither shrink nor expand over time even though methylation levels may change. This is true for 6.5% or 1119 of the 17083 differentially methylated CpG sites (Bonferroni adjusted p < 0.05). For most, the change occurs between birth and age 7 (1041 sites compared to 115 sites between ages 7 and 15-17). Differences that shrink are typically from sites more methylated in males (80% of sites that shrink after birth; 95% of sites that shrink after age 7). For differences that grow, the pattern is not so clear: 84% of sites that grow after birth compared to 25% of sites that grow after age 7 are more methylated in males.

**Fig 5.**
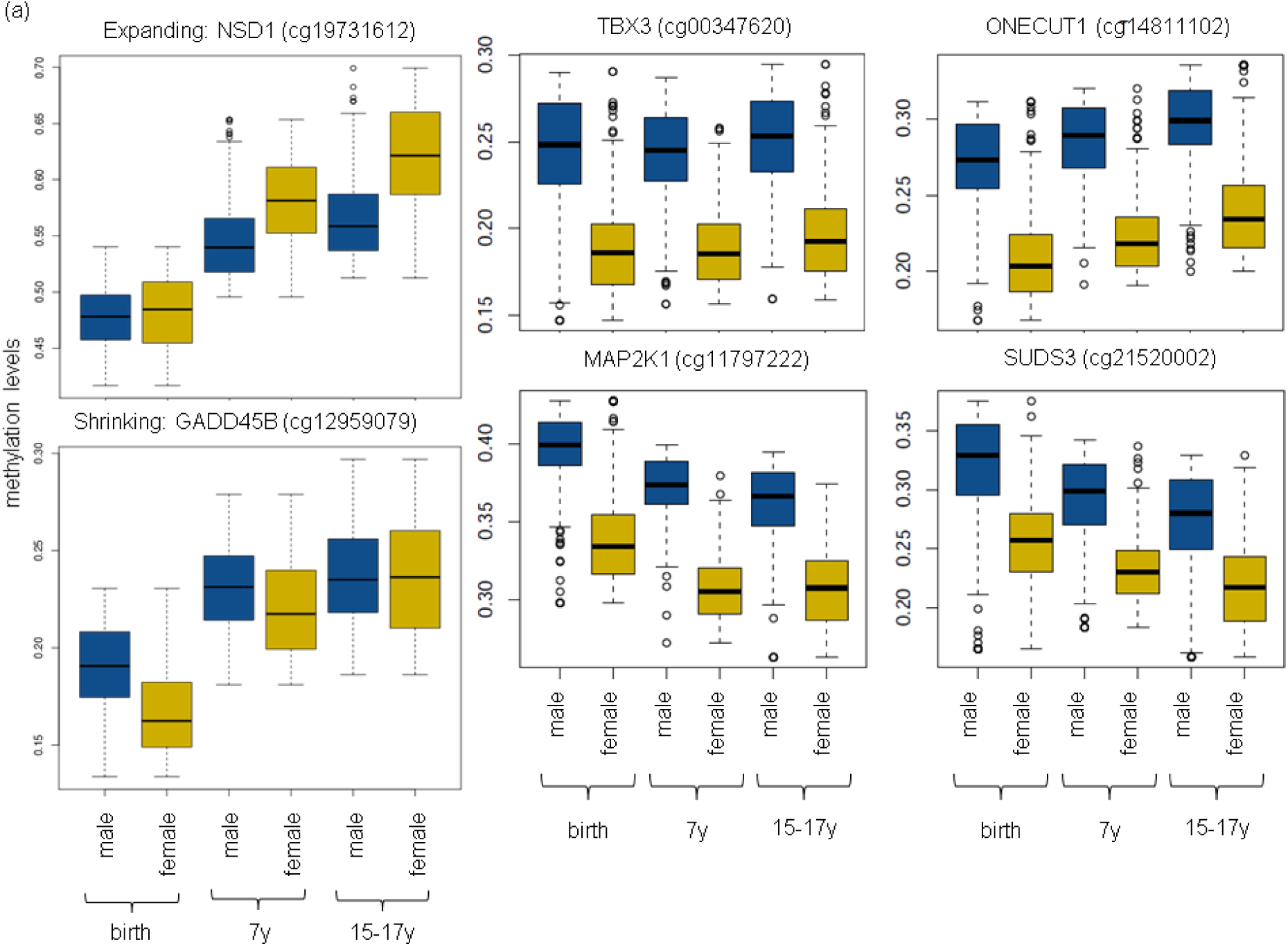
Sex differences and change over time. (a) Examples of sex differences that expand and shrink over time. (b) Examples of sex differences that do not expand or shrink even though methylation levels may or may not change over time.

### Genes with methylation differences that change over time are enriched for genes with developmental functions

CpG sites with methylation that changes over time tend to lie near genes with developmental functions (Spreadsheet S2). Sites with differences that shrink prior to 7y are enriched for genes involved in definitive hemopoiesis, a second wave of hemopoietic stem cell production during early development. Several others may be estrogen targets. A cluster of such sites (p = 0.01, Fisher’s exact test) were observed to change expression in MCF-10F cells following high levels of BPA exposure, and three (p = 0.03) changed following low levels of BPA (19). Eleven of these sites (p = 0.003) appear in regions bound by estrogen receptor following 17β-estradiol treatment in ECC-1 cells (76).

Three sites with differences that shrink after 7y are near the *WT1* gene. *WT1* plays an essential role in development of the urogenital system (77), and following development it is selectively expressed both biallelically and monoallelically in different tissues due to tissue-specific imprinting patterns (78).

Sites with differences that grow prior to 7y are enriched near genes that play roles in neural crest cell migration, pharyngeal system development, peripheral nervous system neuron development, and pattern specification. A differentially methylated gene in each of these processes, *RUNX1,* is possibly more likely regulating the differentiation of hematopoietic stem cells into mature blood cells (79). A second development gene, *HOXD9,* is expressed in cord blood derived mesenchymal stem cells (80), so is also likely important in blood cell differentiation. A third development gene, *HAND2,* is known to play a key role in cardiac morphogenesis as well as limb and branchial arch development (81). These ‘growing’ sites also appear to be enriched near testosterone targets: mouse homologs that are differentially methylated in the bed nucleus of the stria terminalis (BNST) and striatum of adult female mice that had been injected with testosterone shortly after birth (18) (30 BNST genes, p = 2.3×10^−6^; 41 striatum genes, p = 2.5×10^−5^; Fisher’s exact test). Several sites more methylated in females at age 7 and that witness further increases by age 15-17 may be estrogen targets. Five nearby genes (p = 0.03, Fisher’s exact test) have mouse homologs that are differentially methylated in the hippocampus of female mice that had been ovariectomized at 8 weeks (62).

Sites with increased differences after 7y are enriched near the CADPS2 gene and have higher methylation in females. CADPS2 is known to regulate synaptic vesicles and harbours mutations associated with higher risk of autism. Expression of a splice variant of CADPS2 in blood has been linked to intelligence and memory in healthy adults (82).

### Replication of sex-specific methylation

Sex-specific autosomal CpG sites have been identified in earlier Illumina 450K studies for a variety of human tissues. Chen et al. (41) identify autosomal differences but do not report how many remain after cross-hybridizing and polymorphic probes are omitted. Price et al. (35) identify 45 autosomal differences in adult whole blood DNA methylation, Xu et al. (38) 614 in prefrontal cortex, Kaz et al. (40) 82 in colon, Inoshita et al. (36) 292 in peripheral leukocytes, Spiers et al. (39) 521 in human fetal cortex samples spanning 23 to 184 days post-conception, Yousefi et al. (32) 3031 in cord blood, and Singmann et al. (37) 11,010 in peripheral blood from adults aged 32-81y. The sites identified by each except for Kaz et al. have surprisingly large overlaps with our sex-specific sites (p < 2×10^−14^; Table S6). We observed a surprisingly large overlap with fetal cortex (39) with over 180 of their 263 differentially methylated sites differentially methylated at each time point in ARIES. Furthermore, the direction of methylation difference (e.g. higher methylation in males) was conserved for all but three CpG sites: cg26207503 (MYF5 gene) more methylated in males at all time points, cg07173823 (C1orf228 gene) more methylated in males at ages 7 and 15-17, and cg07173823 (KLF13 gene) more methylated in males at age 7.

Several human studies have investigated sex differences in human gene expression (Table S7) (16, 83-87). Unexpectedly, whereas we observe very large overlaps between our differentially methylated genes and genes differentially expressed in brain, we observe little agreement with genes differentially expressed in several other tissues including blood. And although consistently higher methylation levels observed in males would predict higher gene expression levels in females, only one dataset suggests higher female gene expression.

Agreement of our findings with a recent mouse study (88) of gene expression in adipose, muscle, liver and brain tissues is unexpectedly much stronger (Table S8). We observe strong agreement with adipose (p < 1.5×10^−8^; Fisher’s exact test), muscle (p < 1.3×10^−4^) and liver (p < 0.0051) but very little agreement with brain (p = 1). And consistent with our finding of higher methylation in human males, gene expression is generally higher in female mice (51-60% of differentially expressed genes are more expressed in females).

### Replication of overall stability and specific changes over time

To perform more detailed replications, we reanalysed publicly available datasets with DNA methylation profiles obtained from cord (89, 90) and peripheral blood from individuals less than 20y (91-93) (Table S9). For each dataset, agreement is again strong (p < 2.2×10^−5^; Fisher’s exact test), particularly for the larger peripheral blood datasets (p < 2.5×10^−149^) with almost no disagreement about direction of associations (Table 4). Greater prevalence of higher methylation in males however is only observed in one of the two cord and one of the two childhood datasets.

**Table 4.** Sex-specific sites identified in re-analyzed publicly available datasets. The numbers of CpG sites more methylated in males and females for each dataset are provided followed by the numbers replicated in ARIES. CpG sites with inverted association directions are given in the column labelled ‘opposite direction’, and the p-values from Fisher’s exact test of overlap significance are given in the final column.

| GEO accession | More male | More female | Dataset age | ARIES time point | Agree: more in male | Agree: more in female | Opposite | P-value |
| --- | --- | --- | --- | --- | --- | --- | --- | --- |
| GSE62924 (cord; 0y) (89) | 47 | 10 | 0y | birth | 19 | 3 | 0 | $6.8 \times 10^{-12}$ |
| GSE64316 (cord; 0y) (90) | 1 | 3 | 0y | birth | 1 | 3 | 0 | $2.2 \times 10^{-5}$ |
| GSE40576 (6-12y) (93) | 347 | 170 | 6-12y | 7y | 175 | 82 | 7 | $2.5 \times 10^{-149}$ |
| GSE36054 (1-12y) (91) | 62 | 271 | 1-12y | 7y | 47 | 176 | 9 | $4.5 \times 10^{-175}$ |
| GSE56105 (10-19y) (92) | 1184 | 1183 | 10-19y | 15-17y | 622 | 653 | 38 | 0 |

It is not possible to completely test replication of change in sex differences over time because each of the replication datasets contains only cord, childhood or teenage methylation profiles. Consequently, they could not be normalized together to test change across all time points. We were however able to test age-sex interactions prior to 12y and after 10y as well as to check for changes in association direction across 10-12y in different datasets. In the 10-19y dataset (92), four age-sex interactions were identified (Bonferroni adjusted p < 0.05), all indicating increasing methylation differences between males and females over time. Three of these four (cg21184711/CADPS2, cg23256579/PRR4, cg27615582/PRR4) were observed to have age-sex interactions in our data as well, also indicating increasing methylation differences between males and females (Fig S1). In the 6-12y dataset (93), three age-sex interactions were identified (Bonferroni adjusted p < 0.05), all indicating decreasing methylation differences between males and females. None of them, however, replicated findings in ARIES. No age-sex interactions were identified in the 1-12y dataset (91).

No inversions were observed, that is methylation associations that change direction over time, when comparing replication dataset pairs. This replicates the lack of inversions found in our data.

### Replication of greater variability in males

In all replication datasets but one (cord dataset (90)), there were more CpG sites with greater variance in males than there were CpG sites with greater variance in females (double generalized linear model, Bonferroni adjusted p < 0.05). However, the number with greater variance in males relative to females was not nearly as pronounced as in our data (a ratio of 1.1-1.9 in other datasets compared to 3-5 in our data) (Table S10).

### Performance of autosomal sex prediction in replication datasets

Sex predictions in replication datasets were generated by applying k-means clustering (k=2) to the methylation levels of all autosomal CpG sites differentially methylated in ARIES at each time point with at least a 10% methylation difference between males and females. Sex predictions were nearly perfect in each replication dataset. Only a few misclassifications were made: none in the cord datasets (GSE62924 and GSE64316), 8 from 194 in GSE40576, 7 from 465 in GSE56105, and 1 from 118 in GSE36054.

## Discussion

Thousands of profound differences in autosomal cord blood and peripheral blood DNA methylation were observed between males and females in a longitudinal DNA methylation dataset with profiles generated at three time points: birth, 7y and 15-17y. Not only was it possible to observe differences between the sexes, but it was also possible for the first time to observe how these differences changed throughout childhood. The use of Illumina Infinium HM450 BeadChip technology allowed straightforward interpretation of results as well as convenient comparison to a rapidly growing body of publicly available data using the same technology. Since each individual is a member of ALSPAC, we were able to test several hypotheses about relevant phenotypes and exposures using data that has been collected for each individual and their parents over the course of several years.

The study is limited by the coverage of the microarray to only 1.7% of the CpG sites in the human genome, mostly restricted to gene promoter regions. Data about hormone levels in blood were limited to about 20% of the individuals so statistical tests lacked power. DNA methylation profiles were obtained from cord and peripheral blood and some variation in estimated cell type proportions was observed, with the proportions of some cell types confounded with sex. Furthermore, cell types and their proportions in cord blood are somewhat different from those in peripheral blood, and some of the peripheral blood samples came from buffy coats rather than whole blood. However, cell type proportion heterogeneity among the samples appeared to be handled well by including surrogate variables in linear models (94). Including cell type proportions directly as model covariates had very little effect on results. Finally, findings are limited to blood DNA methylation, and DNA methylation is known to differ quite dramatically between tissues. We do however have extremely large overlaps with sex differences found in brain (cortex) in both human fetuses and adults.

Our findings are consistent with the hypothesis that wide-spread sex-discordant autosomal DNA methylation is established very early in fetal development, likely as a response to the presence of male sex hormones, and then stably maintained throughout childhood in spite of dramatic fluctuations in the levels of the same hormones that are associated with the methylation differences in the first place. This stable maintenance results in 70-80% of methylation differences being conserved from one time point to the next and in only 1 in 15 methylation differences changing over time.

Early establishment of sex-specific methylation is supported by the fact that differentially methylated genes are enriched mainly for developmental processes such as organ morphogenesis and sexual development. Furthermore, widespread sex-specific methylation differences identified in fetal brain are almost all found differentially methylated later in cord and peripheral blood at 7y and 15-17y.

Several lines of evidence support the establishment of sex-specific methylation largely as a result of male sex hormone exposure very early in development.

Firstly, given that male sex hormone exposure is known to vary between individuals, dependence of methylation differences on such a variable exposure would suggest greater methylation variation in males. We observe this not only in ARIES at each time point but also in the replication datasets, though not to the same extent.

Secondly, differentially methylated genes are highly enriched for testosterone targets but not for estrogen targets in spite of the fact that estrogen is suspected to directly interact with DNA methylation machinery (22). This is also consistent with enrichment of differentially methylated genes among those most highly expressed in the adrenal cortex and that this enrichment is stronger than any of the 78 other tissues tested including whole blood. The adrenal cortex is part of the hypothalamus-pituitary-adrenal (HPA) axis and produces most of the stress-mediating glucocorticoids and mineralocorticoids. Stress response is known to be extremely sexually dimorphic (95). In addition, the adrenal cortex also secretes sex steroid hormones progestin, androgen, and estrogen (96). Though the amount of sex hormone is a small fraction of that produced by the body, these small secretions play a critical role in development. During pregnancy, the fetal adrenal gland contributes to maternal estrogen levels by secreting prohormones that are aromatized in the placenta (96). Around the age of 6-7, adrenarche is signalled by increased secretion of adrenal androgens due to the development of the zona reticularis in the adrenal cortex (97). In women, 50% of testosterone is converted from adrenal androgens and, although testosterone levels are about 15 times lower in adult females than in adult males, unusually low testosterone levels are known to have a wide range of effects including reduced sexual desire, vaginal health, wellbeing, bone health and lean body mass (98).

Thirdly, both directly and through the use of methylation-derived ‘sex scores’, we show that sex-associated DNA methylation variation is preserved over time. Our hypothesis would predict that this variation should be associated with developmental sex hormone levels. We do not have measurements from this early time point so we instead tested the association with childhood sex hormone levels. Indeed, we identified nominally significant associations (unadjusted p < 0.05), however these associations failed to survive adjustment for multiple testing. We also tested and observed a few nominal associations with phenotypes linked to male sex hormone levels: gender-typed play behaviour, grip strength and finger 2D:4D ratio. Further work in larger datasets is necessary to evaluate the utility of the ‘sex score’ or variations of it in characterizing human sexuality.

Finally, causal analysis using two-sample Mendelian Randomization provided possible support for a causal effect of testosterone on blood DNA methylation.

Replication of these results in publicly available datasets was strong. Although none of the datasets provided longitudinal measures of DNA methylation, we did replicate a large number of the sex discordant CpG sites including a few sites with changing discordance across puberty. Given the strong bias toward higher methylation levels in males in our data, we were surprised that this bias was not consistently replicated. In fact, one dataset (91) indicated the opposite bias.

In all, our findings suggest that sex-discordant autosomal DNA methylation is widespread throughout the genome, and likely due to the first androgen exposures *in utero.* It is then stably maintained from birth to late adolescence despite dramatic fluctuations in the levels of the very same hormones. It thus represents another example where exposure timing is as critical as the exposure itself.

## Materials and Methods

### Study population and sample acquisition

This study used DNA methylation data generated under the auspices of the Avon Longitudinal Study of Parents and Children (ALSPAC) (42, 43). DNA extracted from cord blood and peripheral blood samples at 7 and 17 years along with a wide range of exposure and phenotypic data were used. DNA methylation analysis and data pre-processing were performed at the University of Bristol as part of the Accessible Resource for Integrated Epigenomic Studies (ARIES) project (http://www.ariesepigenomics.org.uk) (44). Data are available from by request from the Avon Longitudinal Study of Parents and Children Executive Committee (http://www.bristol.ac.uk/alspac/researchers/access/) for researchers who meet the criteria for access to confidential data.

### Ethics Statement

Ethical approval for the ALSPAC study was obtained from the ALSPAC Ethics and Law Committee and the local research ethics committees.

### DNA methylation profile generation

DNA was bisulphite converted using the Zymo EZ DNA Methylation™ kit (Zymo, Irvine, CA). Infinium HumanMethylation450 BeadChips (Illumina, Inc.) were used to measure genome-wide DNA methylation levels at over 485,000 CpG sites. The arrays were scanned using an Illumina iScan, with initial quality review using GenomeStudio. This assay detects methylation of cytosines using two site-specific probes - one to detect the methylated (M) locus and one to detect the unmethylated (U) locus. The ratio of fluorescent signals from the methylated site versus the unmethylated site determines the level of methylation at the locus. The level of methylation is expressed as a “Beta” value (β-value), ranging from 0 (no cytosine methylation) to 1 (complete cytosine methylation).

### Quality control

During the data generation process a wide range of batch variables were recorded in a purpose-built laboratory information management system (LIMS). The LIMS also reported QC metrics from the standard control probes on the 450K BeadChip. Samples failing quality (samples with >20% probes with p-value >= 0.01) were repeated. Samples from all three time points in ARIES were randomized across arrays to minimise the potential for batch effects. As an additional QC step, genotype probes on the 450K BeadChip were compared between samples from the same individual and against SNP-chip data to identify and remove any sample mismatches.

### Methylation profile normalization

Raw β-values were pre-processed using R (version 3.0.1) with background correction and subset quantile normalisation performed using the pipeline described by Touleimat and Tost (99) and implemented in the watermelon R package (100). Finally, to reduce influence of outliers in regression models, normalized β-values were 90%-Winsorized.

### Probe exclusions

In addition to excluding probes annotated to CpG sites on chromosomes X and Y, probes with little variance across all methylation profiles (inter-quartile range < 0.01) were excluded as were probes with non-specific binding or that target polymorphic CpG sites (41) or were otherwise previously identified as biased due to the potential presence of multiple SNPs or indels in the probe binding site or discordance with whole genome bisulfite sequencing (101).

### Cell type heterogeneity

Blood is composed of many cell types and composition ratios can vary over time within a given individual as well as between individuals. DNA methylation differs significantly between blood cell types so it is necessary to adjust for cell count heterogeneity in methylation analyses to avoid potential confounding. Cell counts per individual were estimated from DNA methylation profiles using the method described by Houseman et al. (102) using the ‘estimateCellCounts’ function from the ‘minfi’ R package (103) (Table S11). Cell types included CD8+ T cells, CD4+ T cells, CD56 natural killer cells, CD19 B cells, CD14+ monocytes and granulocytes. Some of these estimates are significantly associated with sex (Table S12).

### Methylation differences between males and females

Sex differences were identified in R using linear models implemented in the package ‘limma’ (104), including as covariates the top 20 independent surrogate variables (ISVs) generated by the package ‘ISVA’ package (94). ISVs are included in order to adjust for variation in the methylation data due to demographic, environmental or technical factors in addition to that associated with sex.

As noted above, estimated cell type proportions were significantly associated with sex, however they were not included in the model as covariates because they were well-represented by ISVs (Table S13). Only the low proportion cell types have low correlations. These have little effect on the sex differences that we identify. A reanalysis of the data including both ISVs and estimated cell type proportions as covariates and identified almost identical associations with sex (Table S14).

### DNA methylation change over time

Changes in DNA methylation over time were analysed using a multilevel model (105) fitted to each of the 17K CpG sites identified as differentially methylated between males and females at any of the three ARIES time points (birth, 7y, and 15-17y). The model includes a random intercept for each child and a linear regression spline term to allow a changes in slope after age 7across the second time point. Random slopes were not used due to insufficient between subject variation in methylation change trajectories over the three measurement occasions. Complete details of the model are provided in the Supplementary Information.

### Imprinted genes and imprinting control regions

Imprinted genes were obtained from the geneimprint website (http://www.geneimprint.com: retrieved 31-May-2013). As an approximation of imprinting control regions (ICRs), we used a set of imprinted differentially methylated regions, some of which have been shown to be ICRs by mouse knockout studies (Table S3 of (65)).

### Methylation variance

Double generalized linear models were applied to methylation data at each time point to simultaneously identify mean and variance differences between the sexes (66). Models were implemented in R using the package ‘dglm’.

### Sex score

Sex scores were calculated at each time point with respect to the linear models used to identify methylation differences between the sexes at each time point. For each individual i at time point j, methylation levels m_ijk_ for each differentially methylated CpG site k ∈ D_j_ were adjusted for the previously computed ISVA components and then multiplied by the linear model sex coefficient β_jk_ for that CpG site. The final sex score S_ij_ was then calculated by taking the sum of these across all differentially methylated CpG sites D:

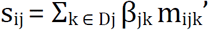

where m_ijk_’ denotes the adjusted methylation level.

### Gene Ontology analysis

Enrichment of Gene Ontology (45) biological processes was calculated by linking processes to each of the non-excluded CpG sites using the R package ‘IlluminaHumanMethylation450k.db’ (106) and enrichment was calculated using a weighted variant of Fisher’s exact test implemented in the topGO R package (107). The topGO algorithm is designed to eliminate local dependencies between GO terms. Statistics are provided in Spreadsheet S2.

### ALSPAC variables

See Supplementary Information.

### Replication datasets and analyses

We considered for replication all publicly available DNA methylation datasets deposited in the Gene Expression Omnibus (http://www.ncbi.nlm.nih.gov/geo/) generated using the Illumina Infinium HumanMethylation450 BeadChip. Eligible datasets needed to be derived from blood samples (cord or peripheral) extracted from healthy individuals under the age of 20 and include at least 10 males and 10 females. Datasets satisfying these criteria are shown in Table S9. Further information about the specifics of each dataset and how they were analysed is given in the Supplementary Information.

## Acknowledgements

We are extremely grateful to all the families who took part in the ALSPAC study, the midwives for their help in recruiting them, and the whole ALSPAC team, which includes interviewers, computer and laboratory technicians, clerical workers, research scientists, volunteers, managers, receptionists and nurses.

The UK Medical Research Council and Wellcome (www.wellcome.ac.uk; 102215/2/13/2) and the University of Bristol provide core support for ALSPAC. This research was specifically funded by the UK Biotechnology and Biological Sciences Research Council (www.bbsrc.ac.uk; BB/I025751/1 and BB/I025263/1); the UK Economic and Social Research Council (www.esrc.ac.uk; ES/N000498/1 to CR); GDS, CR, AS, GS, TG, SR and MS work within the Medical Research Council Integrative Epidemiology Unit at the University of Bristol, which is supported by the UK Medical Research Council (www.mrc.ac.uk; MC_UU_12013/1 and MC_UU_12013/2); the UK Economic and Social Research Council (www.esrc.ac.uk; RES-060-23- 0011 to GDS and CR); and the European Research Council (erc.europa.eu; DEVHEALTH 269874 to GDS).

## Supplementary Information

### Supplementary Materials and Methods

#### DNA methylation change over time

Changes in DNA methylation over time were analysed using a multilevel model (105) fitted to each of the 17K CpG sites identified as differentially methylated between males and females at any of the three ARIES time points (birth, 7y, and 15-17y). The model includes a random intercept for each child and a linear regression spline term to allow a changes in slope after age 7across the second time point. Random slopes were not used due to insufficient between subject variation in methylation change trajectories over the three measurement occasions.

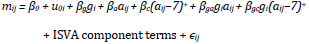

where:

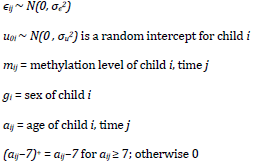

This model implies the following expressions for mean methylation levels:

| Sex ($g_i$) | Time($j$) | Expression | Mean methylation in |
| --- | --- | --- | --- |
| female (0) | 1 | $\beta_0$ | female cord blood |
| male (1) | 1 | $\beta_0 + \beta_g$ | male cord blood |
| female | 2 | $\beta_0 + 7\beta_a$ | female 7y blood |
| male | 2 | $\beta_0 + \beta_g + 7\beta_a + 7\beta_{ga}$ | male 7y blood |
| female | 3 | $\beta_0 + 17\beta_a + 10\beta_c$ | female 17y blood |
| male | 3 | $\beta_0 + \beta_g + 17\beta_a + 17\beta_{ga} + 10\beta_c + 10\beta_{gc}$ | male 17y blood |

As well as the following expression for changes in methylation:

| Expression | Methylation change |
| --- | --- |
| $\beta_a$ | females between birth and 7y |
| $\beta_a + \beta_{ga}$ | males between birth and 7y |
| $\beta_a + \beta_c$ | females after 7y |
| $\beta_a + \beta_{ga} + \beta_c + \beta_{gc}$ | males after 7y |
| $\beta_{ga}$ | difference between sexes up to 7y |
| $\beta_{gc} + \beta_{ga}$ | difference between sexes after 7y |

There is evidence for a methylation sex difference expanding or shrinking over time whenever β_ga_ or β_ga_ + β_gc_ is significantly different from 0.

Note that change in methylation is given as percentage change per year.

#### ALSPAC variables

The Pre-School Activities Inventory (PSAI) is a standardized measure of gender role based on gender-typed play behaviour (108). The PSAI (ALSAPC id: KJ367) was assessed around 42 months of age. The Childhood Activities Inventory (CAI) is an adapted and shortened version of the PSAI. The CAI (ALSPAC id: F8GB041) was assessed around 8.5 years of age.

Sex of older siblings (biological) is not recorded directly in ALSPAC. Instead, several variables identify the sexes and ages of other children living in the same home as the study child (ALSPAC ids: J422-J427). Hence, these data were only used when other data indicated that it was reasonable to assume that the older children in the home were biological siblings and it was unlikely that previous pregnancies were successful. We therefore had to consider only a subset (70%) of study children for whom the following holds (at age 47 months): child lives with their natural mother (J378 = ‘Yes’), all siblings in the household are natural (J382 + J383 + 1 = J428), all older siblings live in the household (J422a + J422b = B005), previous pregnancy child is still alive (B024 = ‘Yes’), no older siblings were stillborn (B012 = 0), and no older siblings died after birth (B014 = 0). Two versions of this variable were considered: the sex of next older sibling and the number of older brothers minus the number of older sisters.

Sex and growth hormones were measured in blood extracted around 8.5 years of age. These included androstenedione (ALSPAC id: andro_bbs), dehydroepiandrosterone (DHEAS; ALSPAC id: dheas_id), growth hormone binding protein (GHBP; ALSPAC id: shbg_bbs), and sex hormone binding globulin (SHBG; ASLPAC id: shgb_bbs). Measurements are typically available only for about 20% of the study members.

Grip strength was assessed using the Jamar hand dynamometer, measured as isometric strength in kilograms. Grip strength here refers to the strength of the dominant hand (ALSPAC ids: FEGS105, FEGS115, FEGS010).

Finger 2D:4D ratio was measured for both left and right hands (ALSPAC ids: FEMS105, FEMS106) from photocopies of participants’ hands using digital calipers at 11y.

Please note that the study website contains details of all the data that is available through a fully searchable data dictionary (http://www.bris.ac.uk/alspac/researchers/data-access/data-dictionary/).

#### Replication datasets and analyses

We considered for replication all publicly available DNA methylation datasets deposited in the Gene Expression Omnibus (http://www.ncbi.nlm.nih.gov/geo/) generated using the Illumina Infinium HumanMethylation450 BeadChip. Eligible datasets needed to be derived from blood samples (cord or peripheral) extracted from healthy individuals under the age of 20 and include at least 10 males and 10 females. Datasets satisfying these criteria are shown in Table S9.

As a quality control step, sex was predicted for each dataset by applying k-means clustering to the normalized methylation estimates on the X and Y chromosomes. The cluster with the higher median X chromosome methylation estimates was identified as female and the other cluster as male. In each case except datasets GSE54399 and GSE40576, predictions agreed perfectly with provided sex designations. For GSE54399, agreement was random so the dataset was excluded from further analysis. For GSE40576, sex information was not provided so predictions from methylation data were used.

For dataset GSE40576, age information was also not provided so we used estimates from methylation data. Estimates were obtained by generating childhood age predictors independently in datasets GSE36054 and GSE56105 and then applying each in the opposite dataset. Pearson correlation with actual age was 0.95 in GSE36054 and 0.83 in GSE56105. Correlation between age estimates obtained from each predictor in GSE40576 was 0.8. Given the fact that GSE56105 is much larger than GSE36054 and the predictor from GSE56105 obtained a much higher correlation with actual age in GSE36054, we elected to use the age estimates from the GSE56105 dataset.

Age predictors were created by first identifying CpG sites significantly associated with age (Bonferroni adjusted p < 0.05). The implementation of elastic net in the R package glmnet (109) was then applied to the DNA methylation levels of these CpG sites in the same dataset to obtain an age predictor.

Methylation differences between males and females were tested as described above for ARIES with two possible exceptions: reduced numbers of ISVA components were included as covariates in the regression model depending on the size of the dataset (maximum 20 or n/20), and for specific datasets one additional phenotype or exposure was included as a covariate.

Autosomal sex predictions were generated by first identifying all CpG sites with at least 10% methylation differences between males and females in ARIES at all three time points. Principal components analysis was then applied to the methylation levels of these CpG sites in the given dataset. K-means clustering was applied to the first two principal components to identify two sample clusters. The most likely sex of the cluster was identified by calculating methylation differences between the two clusters and comparing the signs of the differences to the signs of the effect sizes in ARIES.

## Supplementary Results

### Evidence of genomic clustering

Although differentially methylated sites are distributed throughout the genome, there are surprisingly many on chromosomes 19 and 20 and lesser so on chromosomes 10 and 22 (Bonferroni adjusted p < 0.05; Fisher’s exact test; Table S15).

### Imprinted genes in mouse enriched for testosterone but not estrogen targets

Testosterone targets were identified as genes differentially methylated in the brains of female mice 60 days after being injected with testosterone at birth (18). Imprinted genes were obtained from the geneimprint dataset (http://www.geneimprint.com/). there are 75 imprinted genes in mice. In the bed nucleus of the stria terminalis (BNST), 20 of the 673 differentially methylated genes are imprinted (p < 9×10^−14^, Fisher’s exact test). In the striatum, 16 of the 1251 differentially methylated genes are imprinted (p < 6×10^−6^).

Genes with sex-specific imprinting features in the mouse brain (64) are also enriched with testosterone targets. In the prefrontal cortex, 14 of 40 male-specific paternally expressed genes are testosterone targets in the striatum (p < 3×10^−7^) and, in the hypothalamus, 10 of 38 female-specific maternally expressed genes are testosterone targets in the striatum (p < 5×10^−4^).

Estrogen targets were identified as genes differentially methylated between the hippocampi of ovariectomized mice with and without estradiol treatment (62). Of the 810 estrogen targets, only 4 are imprinted (p > 0.25).

### Males more variably methylated than females

The greater variability in males could be an artifact of males having higher methylation levels. We therefore made repeated random selections of CpG sites M_1_, M_2_, …, M_100_ that are more methylated in males but equal in number and having the same distribution of methylation levels as the sites more methylated in females (we call this latter set F). If greater variance is an artifact of higher methylation levels, then these selections should contain similar numbers of sites more variably methylated in males as F contains sites more variably methylated in females. In fact, we find that F contains less than 3 more variably methylated sites whereas each M_i_ contains 8-15 more variably methylated sites per time point.

## Supplementary Spreadsheets

**Spreadsheet S1**. CpG sites with sex-specific DNA methylation in ARIES.

**Spreadsheet S2**. Gene Ontology biological processes enriched for sex-specific DNA methylation.

---

**Fig S1.**
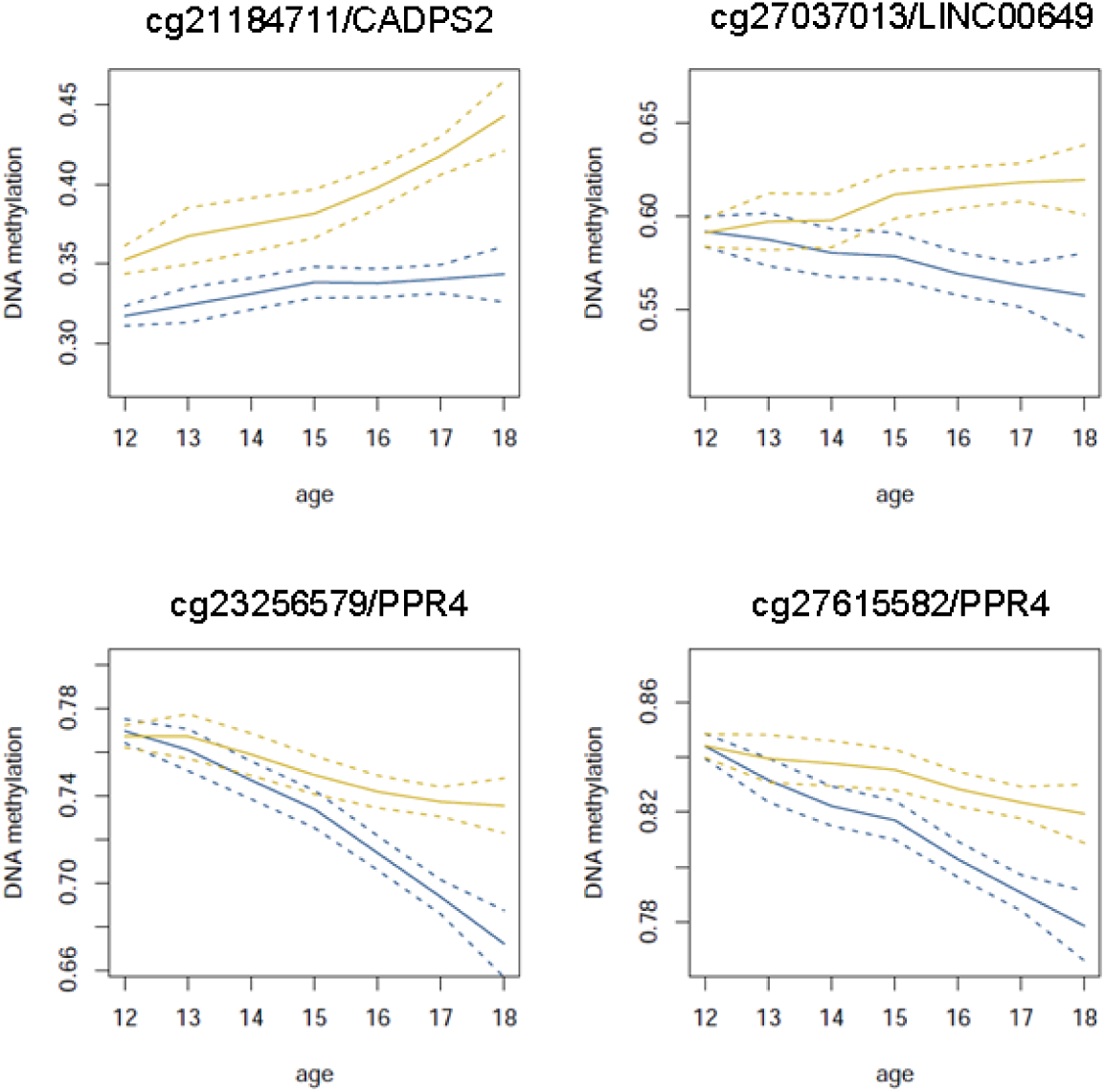
CpG sites with differences that expand over time in the GSE56105 dataset. Solid lines depict Loess-smoothed methylation levels. Dashed lines contain the 95% confidence interval.

**Table S1.** Characteristics of the ARIES sample by sex and measurement age. Characteristics of the ARIES sample by sex and measurement age. Data was collected at birth, age 7, age 15-17. T-tests are used to test numeric differences between males and females and Pearson’s chi-square tests to test differences in proportions.

| ARIES<br>time<br>points | variables | males |  | females |  | p-value |
| --- | --- | --- | --- | --- | --- | --- |
|  |  | mean/n | sd/% | mean | sd/% |  |
| birth | n = 836 | 394 |  | 442 |  |  |
|  | gestational age | 39.5 | 1.5 | 39.6 | 1.5 | 0.16107 |
|  | birthweight | 3552.8 | 503.6 | 3411.5 | 454.6 | 0.00003 |
|  | mother age (months) | 365.3 | 53.1 | 355.4 | 51.8 | 0.00706 |
|  | parity | 0.8 | 0.8 | 0.7 | 0.8 | 0.31067 |
|  | prenatal smoking |  |  |  |  |  |
|  | current | 55 | 14.0 | 62 | 14.0 | 1.00000 |
|  | quit | 16 | 4.1 | 28 | 6.3 | 0.18863 |
|  | never | 293 | 74.4 | 315 | 71.3 | 0.35426 |
|  | caesarean | 36 | 9.1 | 39 | 8.8 | 0.97039 |
| 7y | n = 883 | 437 |  | 446 |  |  |
|  | gestational age | 39.5 | 1.6 | 39.7 | 1.5 | 0.02593 |
|  | birthweight | 3545.9 | 508.7 | 3425.9 | 459.2 | 0.00029 |
|  | mother age (months) | 363.2 | 53.1 | 355.5 | 52.0 | 0.02891 |
|  | parity | 0.7 | 0.8 | 0.7 | 0.9 | 0.71104 |
|  | prenatal smoking |  |  |  |  |  |
|  | current | 61 | 14.0 | 67 | 15.0 | 0.72390 |
|  | quit | 19 | 4.3 | 36 | 8.1 | 0.03156 |
|  | never | 326 | 74.6 | 311 | 69.7 | 0.12396 |
|  | caesarean | 42 | 9.6 | 40 | 9.0 | 0.83143 |
| 15-17y | n = 881 | 426 |  | 455 |  |  |
|  | gestational age | 39.5 | 1.6 | 39.7 | 1.5 | 0.05878 |
|  | birthweight | 3573.6 | 510.4 | 3421.3 | 467.9 | 0.00001 |
|  | mother age (months) | 365.0 | 53.7 | 355.6 | 52.2 | 0.00842 |
|  | parity | 0.8 | 0.8 | 0.7 | 0.8 | 0.25708 |
|  | prenatal smoking |  |  |  |  |  |
|  | current | 56 | 13.1 | 63 | 13.8 | 0.83725 |
|  | quit | 20 | 4.7 | 35 | 7.7 | 0.08943 |
|  | never | 316 | 74.2 | 323 | 71.0 | 0.32495 |
|  | caesarean | 38 | 8.9 | 40 | 8.8 | 1.00000 |
|  | smoking daily (15-17y) | 46 | 10.8 | 53 | 11.6 | 0.76984 |
| all time points | n = 700 | 335 |  | 365 |  |  |
|  | gestational age | 39.5 | 1.5 | 39.7 | 1.5 | 0.07704 |
|  | birthweight | 3569.1 | 502.9 | 3409.8 | 459.8 | 0.00002 |
|  | mother age (months) | 364.9 | 54.6 | 355.1 | 52.4 | 0.01608 |
|  | parity | 0.8 | 0.8 | 0.7 | 0.8 | 0.37894 |
|  | prenatal smoking |  |  |  |  |  |
|  | current | 47 | 14.0 | 51 | 14.0 | 1.00000 |
|  | quit | 15 | 4.5 | 27 | 7.4 | 0.14277 |
|  | never | 248 | 74.0 | 259 | 71.0 | 0.41017 |
|  | caesarean | 29 | 8.7 | 32 | 8.8 | 1.00000 |
|  | smoking daily (15-17y) | 35 | 10.4 | 44 | 12.1 | 0.58116 |

**Table S2.** Sexual developmental genes linked to CpG sites with sex-specific DNA methylation. Numbers of CpG sites more methylated in males (‘M’) and females (‘F’) are given for each ARIES time point (birth, age 7y and age 15-17y). Most of genes were identified in mouse studies so those without human homologs are noted. Genes on the sex chromosomes were not analyzed in our study. These are also noted. For each function, p-values from Fisher’s exact test denote enrichment of sex-specific methylation among CpG sites linked to the genes.

|  |  |  | birth |  | 7y |  | 15-17y |  |
| --- | --- | --- | --- | --- | --- | --- | --- | --- |
| function | gene | chr | M | F | M | F | M | F |
| DHEA-S (110) | CYP11A1 | 15 |  |  |  |  |  |  |
|  | CYP17A1 | 10 |  |  |  |  |  |  |
|  | MC2R | 18 |  |  |  |  |  |  |
|  | POR | 7 | 1 |  | 1 |  | 2 |  |
|  | STAR | 8 |  |  |  |  |  |  |
|  | SULT2A1 | 19 |  |  | 1 |  | 1 |  |
|  | p (Fisher's exact test) |  | 1 |  | 0.9 |  | 0.6 |  |
| Genitalia development (111) | ATF3 | 1 | 1 |  |  |  |  |  |
|  | FGF8 | 10 | 2 |  | 3 |  | 3 |  |
|  | GATA4 | 8 | 12 |  | 19 |  | 19 |  |
|  | NR0B1 | X | sex chromosome |  |  |  |  |  |
|  | SHH | 7 | 3 | 2 | 2 | 2 | 2 | 2 |
|  | SOX9 | 17 | 2 |  | 4 |  | 2 |  |
|  | SRY | Y | sex chromosome |  |  |  |  |  |
|  | WNT4 | 1 | 1 |  | 1 |  | 1 |  |
|  | WNT5A | 3 | 6 |  | 6 |  | 6 |  |
|  | ZFPM2 | 8 |  |  |  |  |  |  |
|  | p (Fisher's exact test) |  | 4.8x10<sup>-4</sup> |  | 6.8x10<sup>-6</sup> |  | 3.7x10<sup>-6</sup> |  |
| HOX:Early gonadal development (112) | EMX2 | 10 | 4 |  | 7 |  | 6 |  |
|  | LHX1 | 17 | 8 |  | 9 |  | 9 |  |
|  | LHX9 | 1 | 4 |  | 4 |  | 3 |  |
|  | p (Fisher's exact test) |  | 7x10<sup>-4</sup> |  | 6.8x10<sup>-6</sup> |  | 3.67x10<sup>-6</sup> |  |
| HOX:Oogenesis (112) | CPHX1 |  | mouse gene without human homolog |  |  |  |  |  |
|  | IRX3 | 16 | 8 |  | 11 |  | 12 |  |
|  | LHX8 | 1 | 1 | 8 | 1 | 9 | 2 | 8 |
|  | NOBOX | 7 |  |  |  |  |  |  |
|  | OBOX1 |  | mouse gene without human homolog |  |  |  |  |  |
|  | OBOX6 |  | mouse gene without human homolog |  |  |  |  |  |
|  | p (Fisher's exact test) |  | 1.2x10<sup>-5</sup> |  | 6.5x10<sup>-9</sup> |  | 2.4x10<sup>-7</sup> |  |
| HOX:Sex differentiation and development (112) | ARX | X | sex chromosome |  |  |  |  |  |
|  | ESX1 | X | sex chromosome |  |  |  |  |  |
|  | HOXA11 | 7 | 3 |  | 4 |  | 5 |  |
|  | HOXA13 | 7 | 2 |  | 5 | 1 | 5 |  |
|  | HOXA7 | 7 | 3 |  | 2 |  | 2 |  |
|  | HOXD13 | 2 |  |  |  |  |  |  |
|  | LBX2 | 2 | 2 |  | 3 | 1 | 5 | 1 |
|  | MKX | 10 | 1 |  | 2 |  | 2 | 1 |
|  | PBX1 | 1 | 2 |  | 1 |  |  |  |
|  | RHOX6 |  | mouse gene without human homolog |  |  |  |  |  |
|  | RHOX9 |  | mouse gene without human homolog |  |  |  |  |  |
|  | **p (Fisher's exact test)** |  | 0.3 |  | 0.035 |  | 0.0027 |  |
| HOX:Spermato<br>genesis (112) | PBX4 | 19 |  | 2 |  | 2 |  | 2 |
|  | POU5F2 | 5 |  |  |  |  |  |  |
|  | RHOX5 |  | mouse gene without human homolog |  |  |  |  |  |
|  | TGIF1 | 18 | 1 |  | 1 | 1 | 1 |  |
|  | **p (Fisher's exact test)** |  | 0.3 |  | 0.1 |  | 0.3 |  |

**Table S3.** Correlation of sex-discordant methylation over time. Median correlation (Spearman’s Rho) of methylation levels at sex-discordant CpG sites across time points in males and the median difference between males and females. The p-value indicates the strength of the difference correlation difference between males and females (Wilcoxon rank-sum test).

| | median $R_{\text{male}}$ | median<br>($R_{\text{male}} - R_{\text{female}}$) | P-value |
| --- | --- | --- | --- |
| birth to 7y | 0.13 | 0.0002 | 0.86 |
| birth to 15-17y | 0.11 | -0.004 | $1.2 \times 10^{-11}$ |
| 7y to 15-17y | 0.17 | -0.01 | $3.3 \times 10^{-77}$ |

**Table S4.** Causal analysis of testosterone on sex-specific methylation. Statistics for the two-sample Mendelian randomization analysis of testosterone levels on the DNA methylation sex score.

|  |  |  | Direct genotype associations with testosterone (70) |  | Direct genotype associations with sex score in ARIES (males; all) |  | Wald ratio estimate |  |
| --- | --- | --- | --- | --- | --- | --- | --- | --- |
| Instrumental variable (SNP) | Effect allele | Effect allele frequency | N | $\beta$ (SE) | N | $\beta$ (SE) | $\beta$ (SE) | p-value |
| rs4149056 | T | 0.82 | 21791 | 0.029 (0.0051) | 307; 639 | -0.77 (0.24); -0.42 (0.18) | -26.7 (8.4); -14.3 (6.15) | 0.0015; 0.0199 |

**Table S5.**
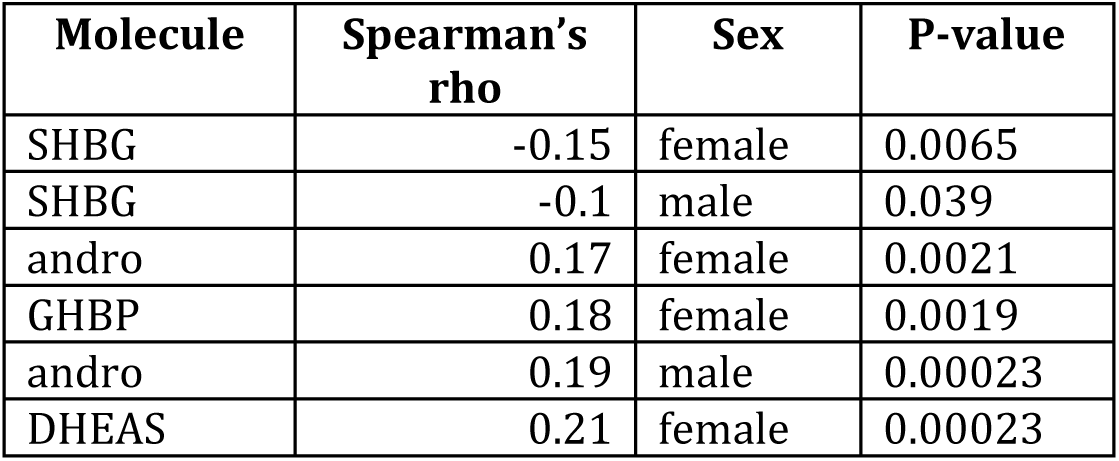
Associations between grip strength and sex hormones. Associations (unadjusted p < 0.05) between grip strength (dominant hand) and molecular measurements taken from peripheral blood at age 8.5y. Molecules tested were androstenedione (andro), dehydroepiandrosterone (DHEAS), growth hormone binding protein (GHBP), sex hormone binding globulin (SHBG).

| Molecule | Spearman's rho | Sex | P-value |
| --- | --- | --- | --- |
| SHBG | -0.15 | female | 0.0065 |
| SHBG | -0.1 | male | 0.039 |
| andro | 0.17 | female | 0.0021 |
| GHBP | 0.18 | female | 0.0019 |
| andro | 0.19 | male | 0.00023 |
| DHEAS | 0.21 | female | 0.00023 |

**Table S6.** Overlap with previously identified sex differences. The table shows the overlap between sex-specific autosomal sites identified in previous Illumina 450K studies and those identified in ARIES. The number of autosomal sites is provided for each study and next to it the number of these sites retained in ARIES after all probe exclusions (see Supplementary Materials and Methods). The non-excluded number is typically much smaller due to our exclusion of potentially biased sites identified by Naeem et al. (101). Most studies excluded sites identified by Chen et al. (41) only. P-values for each time were obtained by Fisher’s exact test with Bonferroni-adjustment for multiple tests.

| Study | Tissue | Autosomal sites | Non-excluded | Cord (11966 sites) | 7y (13672 sites) | 15-17y (12215 sites) |
| --- | --- | --- | --- | --- | --- | --- |
| Price et al. (35) | adult whole blood | 18 | 18 | 2.2x10<sup>-15</sup> (15) | 1.6x10<sup>-14</sup> (15) | 4.2x10<sup>-17</sup> (16) |
| Xu et al. (38) | adult prefrontal cortex | 614 | 250 | 2.6x10<sup>-64</sup> (114) | 1.8x10<sup>-51</sup> (107) | 3.4x10<sup>-53</sup> (104) |
| Kaz et al. (40) | adult colon | 82 | 4 | 0.2 (1) | 0.3 (1) | 0.3 (1) |
| Spiers et al. (39) | fetal cortex | 521 | 263 | 2.3x10<sup>-163</sup> (193) | 6.1x10<sup>-145</sup> (188) | 7.3x10<sup>-148</sup> (184) |
| Inoshita et al. (36) | adult peripheral leukocytes | 292 | 150 | 7.7x10<sup>-69</sup> (93) | 2.3x10<sup>-74</sup> (101) | 1.7x10<sup>-73</sup> (97) |
| Yousefi et al. (32) | cord blood | 3,031 | 1,662 | 10<sup>-300</sup> (838) | 10<sup>-300</sup> (903) | 10<sup>-300</sup> (828) |
| Singmann et al. (37) | adult whole blood | 11,010 | 6,107 | 10<sup>-300</sup> (1934) | 10<sup>-300</sup> (2524) | 10<sup>-300</sup> (2369) |

**Table S7.** Significance of agreement with previous gene expression studies. The study tissue and number of genes identified as differentially expressed between males and females along with the number that are autosomal is shown. The percentage of genes more expressed in females is given in the column denote ‘%’. The column ‘P-value’ provides the statistical significance of the overlap between the autosomal genes differentially expressed in the study and the differentially methylated genes in ARIES (Fisher’s exact test).

| Study | Tissue | Genes | Autosomal | % | P-value |
| --- | --- | --- | --- | --- | --- |
| Reinius et al. (16) | occipital cortex | 934 | 897 | 8 | 6.1x10<sup>-16</sup> |
| Mayne et al. (83) | anterior cingulate cortex | 1616 | 1616 | 35 | 1x10<sup>-12</sup> |
| Mayne et al. (83) | nucleus accumbens | 215 | 215 | 75 | 3.3x10<sup>-7</sup> |
| Mayne et al. (83) | hippocampus | 166 | 166 | 65 | 3.7x10<sup>-7</sup> |
| Mayne et al. (83) | dorsolateral prefrontal cortex | 161 | 161 | 69 | 5.61x10<sup>-6</sup> |
| Mayne et al. (83) | heart | 283 | 283 | 60 | 6.9x10<sup>-6</sup> |
| Mayne et al. (83) | frontal cortex | 8 | 8 | 57 | 0.011 |
| Mayne et al. (83) | colon | 156 | 156 | 26 | 0.012 |
| Simon et al. (84) | platelet RNA | 53 | 44 | 23 | 0.03 |
| Mayne et al. (83) | lung | 3 | 3 | 100 | 0.11 |
| Xu et al. (85) | peripheral blood | 105 | 86 | - | 0.12 |
| Mayne et al. (83) | cerebellum | 40 | 40 | 58 | 0.12 |
| Whitney et al. (86) | peripheral blood | 32 | 24 | 73 | 0.13 |
| Mayne et al. (83) | thyroid | 126 | 126 | 25 | 0.15 |
| Eady et al. (87) | PBMC | 28 | 19 | 40 | 0.33 |
| Mayne et al. (83) | kidney | 177 | 177 | 28 | 0.42 |
| Mayne et al. (83) | liver | 16 | 16 | 75 | 0.50 |

**Table S8.** Replication of sex differences in mice. Size of overlap between genes differentially expressed in mice (88) and those differentially methylated in our study between males and females. P-values were obtained by Fisher’s exact test with Bonferroni-adjustment for multiple tests.

| Mouse tissue | Time point | Mouse genes* | Human genes+ | Overlap | P-value |
| --- | --- | --- | --- | --- | --- |
| adipose | birth | 4641 | 5554 | 1724 | 2.1x10<sup>-6</sup> |
|  | 7y | 4641 | 5777 | 1813 | 1.5x10<sup>-8</sup> |
|  | 15-17y | 4641 | 5357 | 1685 | 4.8x10<sup>-8</sup> |
| brain | birth | 292 | 5554 | 100 | 1 |
|  | 7y | 292 | 5777 | 101 | 1 |
|  | 15-17y | 292 | 5357 | 87 | 1 |
| liver | birth | 3780 | 5554 | 1376 | 0.0051 |
|  | 7y | 3780 | 5777 | 1412 | 0.0416 |
|  | 15-17y | 3780 | 5357 | 1299 | 0.2 |
| muscle | birth | 1762 | 5554 | 665 | 0.005 |
|  | 7y | 1762 | 5777 | 707 | 1.3x10<sup>-4</sup> |
|  | 15-17y | 1762 | 5357 | 658 | 2.2x10<sup>-4</sup> |

\* Number of differentially expressed mouse genes with human homologs. + Number of human genes with mouse homologs and Illumina 450k probes.

**Table S9.** Re-analyzed replication datasets. Each dataset contains at least 10 Illumina Infinium^®^ HM450 BeadChip methylation profiles per sex obtained either from cord or peripheral blood DNA for healthy individuals 0-20y. A subset of GSE36054 including only individuals age 1-12 was analyzed. This excluded only 16 individuals age 13-17. Dataset GSE56105 was similarly restricted to individuals less than age 20.

| GEO accession and ref | cell type | age (mean) | n | male | female | other variables |
| --- | --- | --- | --- | --- | --- | --- |
| GSE54399* | cord blood | 0 | 24 | 14 | 10 |  |
| GSE62924 (89) | cord blood | 0 | 38 | 16 | 22 | arsenic |
| GSE64316 (90) | cord blood | 0 | 20 | 10 | 10 | prenatal tobacco exposure |
| GSE40576 (93) | peripheral blood mononuclear cells | 6-12 (9) | 194 | 97 | 97 | asthma |
| GSE36054 (91) | peripheral blood leukocytes | 1-12 (3.3) | 118 | 79 | 55 |  |
| GSE56105 (92) | peripheral blood lymphocytes | 10-19 (13.8) | 465 | 242 | 223 |  |

\* Dataset GSE54399 was excluded from replication because author sex designations disagreed with sex prediction based on probes targeting chromosomes X and Y.

**Table S10.** Replication of sex-specific variance. Number of CpG sites differentially methylated between males and females in replication datasets that are significantly more variable in males than in females, and vice versa.

| GEO accession and ref | cell type | age (mean) | males more variable | females more variable |
| --- | --- | --- | --- | --- |
| GSE62924 (89) | cord blood | 0 | 0 | 1 |
| GSE64316 (90) | cord blood | 0 | 1 | 0 |
| GSE40576 (93) | peripheral blood mononuclear cells | 6-12 (9) | 50 | 26 |
| GSE36054 (91) | peripheral blood leukocytes | 1-12 (3.3) | 54 | 39 |
| GSE56105 (92) | peripheral blood lymphocytes | 10-19 (13.8) | 59 | 54 |

**Table S11.** Cell counts in ARIES. Median cell type proportions per time point estimated from DNA methylation profiles.

| cell type | birth | 7y | 15-17y |
| --- | --- | --- | --- |
| Granulocytes | 0.52 | 0.48 | 0.52 |
| CD4+ T cells | 0.11 | 0.15 | 0.13 |
| CD8+ T cells | 0.12 | 0.16 | 0.15 |
| CD19+ B cells | 0.15 | 0.14 | 0.11 |
| CD14+ Monocytes | 0.1 | 0.07 | 0.06 |
| CD56+ Natural killer cells | 0.04 | 0.01 | 0.03 |

**Table S12.** Sex-specific cell counts in ARIES. Mean differences and significances (Wilcoxon rank-sum test) of the differences of cell type proportions between males and females.

| cell type | birth |  | 7y |  | 15-17y |  |
| --- | --- | --- | --- | --- | --- | --- |
|  | mean<br>M - F | p-value | mean<br>M - F | p-value | mean<br>M - F | p-value |
| Granulocytes | -0.017 | 0.0042 | -0.004 | 0.5 | -0.02 | 0.0012 |
| CD4+ T cells | 0.008 | 0.0015 | 0.001 | 0.8 | -0.001 | 0.9 |
| CD8+ T cells | -0.002 | 0.2 | 0.003 | 0.4 | -0.005 | 0.1 |
| CD19+ B cells | 0.008 | 0.0076 | 0.006 | $9 \times 10^{-4}$ | 0.012 | $2.6 \times 10^{-7}$ |
| CD14+ Monocytes | 0 | 0.6 | 0.001 | 0.2 | 0.003 | 0.052 |
| CD56+ Natural killer cells | 0.005 | 0.031 | -0.003 | 0.0051 | 0.011 | $3 \times 10^{-5}$ |

**Table S13.** Cell count variation in surrogate variables. Correlation coefficients (Spearman’s Rho) were calculated between estimated cell type proportions and ISVA components. Shown are the largest correlation coefficients (absolute value) for each cell type.

| cell types | birth | 7y | 17y |
| --- | --- | --- | --- |
| Granulocytes | 0.90 | 0.95 | 0.95 |
| CD4+ T cells | 0.81 | 0.83 | 0.80 |
| CD8+ T cells | 0.60 | 0.70 | 0.67 |
| CD19+ B cells | 0.53 | 0.65 | 0.66 |
| CD14+ Monocytes | 0.27 | 0.20 | 0.39 |
| CD56+ Natural killer cells | 0.43 | 0.27 | 0.46 |

**Table S14.** Cell count sensitivity results. Sites differentially methylated under the original model (ISVA components as covariates) and under the new model (ISVA and estimated cell type proportions).

|  | birth | F7 | 15up |
| --- | --- | --- | --- |
| original (ISVA) | 11966 | 13672 | 12215 |
| new (ISVA + cell types) | 11848 | 13412 | 11555 |
| overlap | 11631 | 13137 | 11178 |

**Table S15.** Chromosomal enrichment of sex-specific methylation. Chromosomes with enrichment for sex-specific DNA methylation. Shown are Bonferroni adjusted p-values from Fisher’s exact test.

| Chromosome | Female<br>cord | Female<br>7y | Female<br>15-17y | Male<br>cord | Male<br>7y | Male<br>15-17y |
| --- | --- | --- | --- | --- | --- | --- |
| chr19 | $2.2 \times 10^{-44}$ | $1.3 \times 10^{-31}$ | $1.5 \times 10^{-29}$ | 0.013 | 0.099 | 0.0067 |
| chr20 | $3.3 \times 10^{-16}$ | $7.6 \times 10^{-14}$ | $4.2 \times 10^{-13}$ | 1.0 | 1 | 1 |
| chr22 | 0.0018 | 0.008 | 0.063 | 0.6 | 1 | 1 |
| chr10 | 1 | 0.075 | 0.035 | 1.0 | 1 | 1 |

